# Ribosome-linked mRNA-rRNA chimeras reveal active novel virus host associations

**DOI:** 10.1101/2020.10.30.332502

**Authors:** J. Cesar Ignacio-Espinoza, Sarah M. Laperriere, Yi-Chun Yeh, Jake Weissman, Shengwei Hou, Andrew M. Long, Jed A. Fuhrman

**Author notes:** Department of Oceanography and Coastal Sciences, Louisiana State University, Baton Rouge, LA 70803. Correspondence to be addressed to &.

## Abstract

Viruses of prokaryotes greatly outnumber their hosts^1^ and impact microbial processes across scales, including community assembly, evolution, and metabolism^1^. Metagenomic discovery of novel viruses has greatly expanded viral sequence databases, but only rarely can viral sequences be linked to specific hosts. Here, we adapt proximity ligation methods to ligate ribosomal RNA to transcripts, including viral ones, during translation. We sequenced the resulting chimeras, directly linking marine viral gene expression to specific hosts by transcript association with rRNA sequences. With a sample from the San Pedro Ocean Time-series (SPOT), we found viral-host links to Cyanobacteria, SAR11, SAR116, SAR86, OM75, and Rhodobacteracae hosts, some being the first viruses reported for these groups. We used the SPOT viral and cellular DNA database to track abundances of multiple virus-host pairs monthly over 5 years, e.g. with *Roseovarius* phages tracking the host. Because the vast majority of proximity ligations should occur between an organism’s ribosomes and its own transcripts, we validated our method by looking for self- vs non-self mRNA-rRNA chimeras, by read recruitment to marine single amplified genomes; verifiable non-self chimeras, suggesting off-target linkages, were very rare, indicating host-virus hits were very unlikely to occur by mistake. This approach in practice could link any transcript and its associated processes to specific microorganisms.

---

Microbial communities are central players in the elemental transformations that sustain life on our planet^2^. Like all of life on earth, microbes are susceptible to viral infections, and in the ocean, viruses dominate and exceed their hosts’ numbers manifold^1^. While the importance of virus-host interactions is well recognized, the vast majority viruses observed microscopically and those discovered in recent years by metagenomic approaches do not have a specifically known host, so who infects whom remains largely unknown. This problem is nearly axiomatic in non-marine environments as well, with potential economic, environmental, and health related impacts. Virus-hosts links are therefore one of the main open questions in environmental microbiology, and multiple methods have been developed to address this question, ranging from the informatic (e.g. finding similarities between virus and host genomic sequences^3^) to recent wet-lab based protocols such as viral tagging^4^, finding viruses in single amplified genomes^5^ or adsorbed to cells^6^, digital PCR^7^ and DNA-DNA proximity ligation^8,9^.

Viruses function and reproduce only when their genes are transcribed and then translated to proteins by the host’s ribosomal machinery. We adapted and applied the well-established proximity ligation approach^10^, previously used to probe DNA^9^, RNA^11^ and Protein^12^ interactions, to bond the viral transcripts to the host’s ribosomal RNA after chemically fixing them both in the act of translation, thus allowing us to associate viral gene transcripts directly to the host that is translating them. The bioinformatic analysis allowing us to interpret these connections as virus-host associations is only possible because we now have large libraries of known marine viral sequences determined metagenomically^13,14^ as well as a massive database of ribosomal RNA sequence from organisms across the entire phylogenetic tree^15^. Chimeric mRNA-rRNA sequences that have a virus gene fragment next to a host ribosomal RNA sequence fragment are a “smoking gun” to show an active virus-host link, contrasting with other recent methods that also report more incidental relationships. Here we present proof of concept and the first field application of rRNA-mRNA proximity ligation to specifically link viruses and their hosts through this viral-transcript-host-ribosome connection. We also validate the method by showing that among rRNA-mRNA chimeras, the vast majority are coded by the same organism’s genome, suggesting false, cross-organism, linkages are exceedingly rare.

## Summary of Methods

We present XRM-Seq (Ribosome cross-linking and sequencing). Briefly (Figure 1a) described (See extended methods below): Seawater samples collected in February of 2020 at the San Pedro Ocean Time series station (SPOT^16^) were filtered serially through an 80 um nylon mesh and a 1.2 um fiber glass filter to remove most eukaryotes; cells in the filtrate, containing mostly free-living prokaryotes, were collected on a 0.2 μm filter. (1) These cells were fixed with 1% formaldehyde, cross-linking adjacent proteins to hold ribosomes together. (2) Intact crosslinked ribosomes and total RNA were extracted using acid phenol-chloroform, particularly from the interphase^12^, where the protein-RNA complexes migrate. DNA was depleted with two rounds of DNAse treatment. (3) RNA was randomly cut with micrococcal S1 nuclease, cleaving accessible rRNA strands in the ribosome (Suppl. Fig. 1), which is held together by covalent bonds from crosslinking between the proteins that make most of the ribosomal mass, and it also generates free ends in the mRNA. (4) Free RNA ends (from ribosomal or messenger RNA) were then ligated into circular forms with a circRNA ligase^11^. (5) To enrich for chimeric reads, samples were then subject to degradation of non-circular RNA using RNAse R^11^. Crosslinks were cut by proteinase K. (6) RNA was then retrotranscribed to cDNA, which was then used to prepare libraries that were then deeply sequenced by Illumina Nova Seq. Merged and QC’ed reads (see extended methods below) were then searched for chimeric reads by aligning them against Silva v 132^15^, single cell genomes^17^, a local rRNA sequence database^18^, a set of local viral contigs (representing 5 years of monthly samples) from our previous metagenomic work^13^ and our in-house database of fully sequenced marine viruses, fosmids and assemblies from global marine virus metagenomics projects.

**Figure 1.**
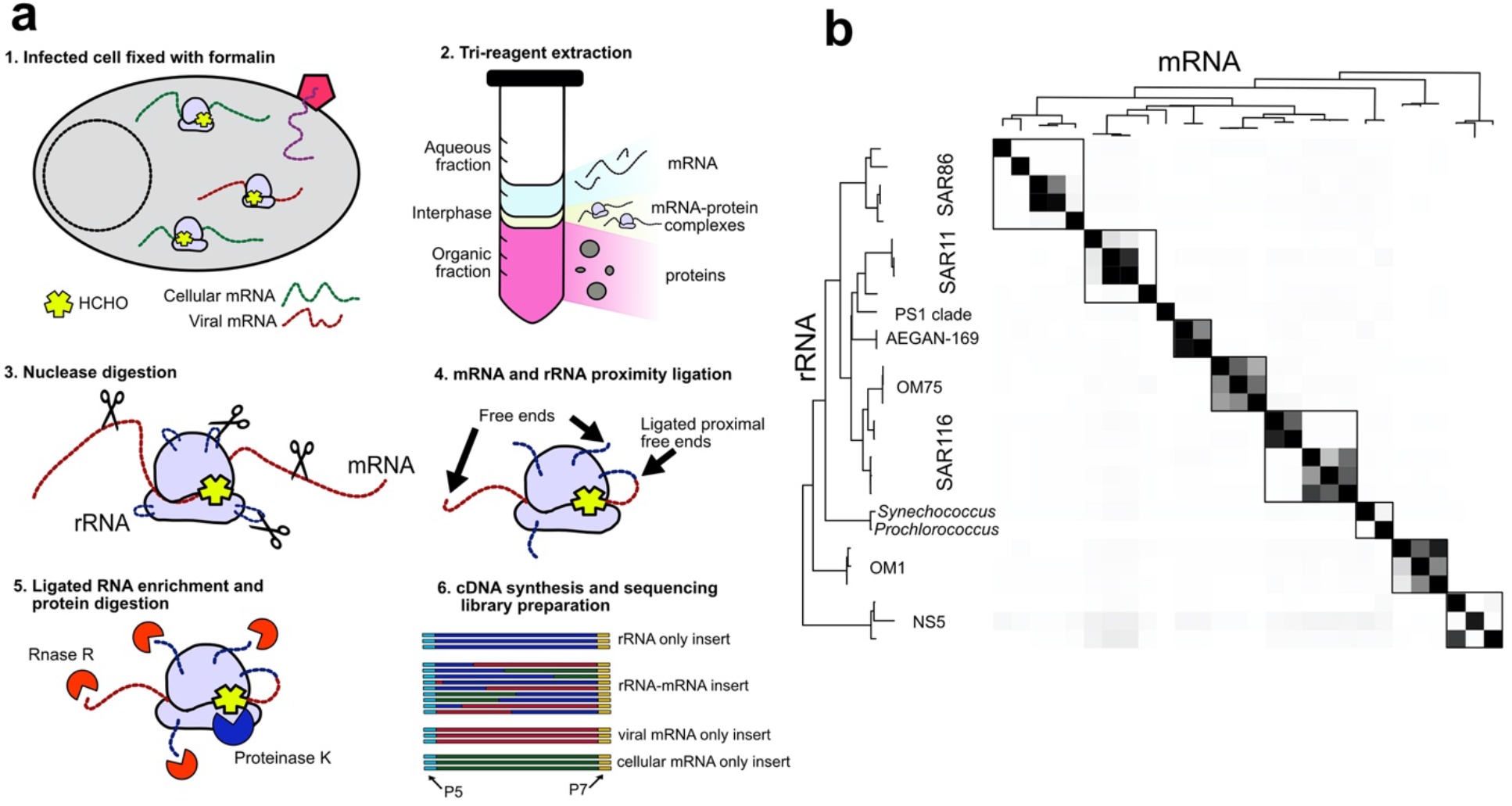
**a)** Diagram of the general method: (1) *In vivo* cross-linking of infected cells in which formalin fixes ribosome (light purple) complexes during translation of host (green) or virus (red) mRNA. (2) Acid phenol chloroform extraction separates the components of the cell lysate; ribosome-mRNA complexes migrate to the interphase. (3) Ribosome complexes are subject to a nuclease digestion that generates free ends in the rRNA (blue) and mRNA (red). (4) Free proximal ends are ligated, which generates chimeras. (5) To enrich for ligated RNA, RNA with exposed ends is degraded with RNAse R (orange pie); Crosslinking is reversed with Proteinase K (dark blue pie). (6) Sequencing libraries were prepared from retrotranscribed purified RNA, many where chimeric, containing host rRNA (Blue) and viral (red) or host (green) mRNA. **b)** Validation, based upon mRNA-rRNA chimeras within SAGs, is demonstrated by the strong domination of within-organism linkages, as expected if the method only links mRNA and rRNA within each cell, not between cells. This is shown by the taxonomic distribution of chimerically-linked ribosomal RNA (y-axis) and mRNA transcripts (x-axis), as determined by sequences mapping to a subtropical-tropical single amplified genome dataset representing all major marine prokaryotic lineages. Intensity is normalized within rows, reflecting number of linkages, as determined by sequences mapping (>95 %ID) to a subtropical-tropical single amplified genome dataset. Tree on left and top (mirrored) is based on 16S rRNA sequences, not similarity among rows/columns. Note the vast majority of all linkages are either to the identical SAG or to a very closely related one, the latter probably reflecting situations where our local organism had no identical SAG but two different very close relatives in the SAG database (accuracy, i.e. the fraction of self-hits within the shown boxes, is 95%, with an associated *p*-value <2.2×10^−16).

## Results and Discussion

### We validated our method and found a negligible rate of false positive linkages

The general validity of the approach was verified by examining chimeric linkages for those between rRNA and protein-coding genes from the same organisms, which would be expected to be the vast majority of linkages if the method worked as planned. For this validation we wanted to use independently-determined and bona fide genome sequences, so we did not try to bin genomes from our own local metagenomic data (to avoid assembly artifacts or any possible circular reasoning), but instead used published marine genomes as a reference set. In particular, we used sequences from a recently released set of tropical and subtropical marine single amplified genomes (SAGs)^17^. This analysis showed that the vast majority these linkages were to the same SAG or within the same close lineage (Figure 1b); Because these SAGs were not from our exact site, we expect that these close (but not perfect) hits are due to database imperfections (i.e. our local organism may not match a SAG exactly but may have two similarly-close SAG relatives) and not because erroneous, i.e. non-self, linkages somehow always happened to occur only with close relatives. Erroneous linkages would instead be expected to occur randomly among and across the several most abundant and diverse taxa, which was not observed. Thus these results validate our approach, and we felt confident extending the analyses to virus-host linkages

### mRNA-rRNA chimeras reveal novel virus-host links

To cast the widest possible net for potential viruses and hosts in the chimeric linkages, we took advantage of the fact that we have been studying the SPOT site for several years, and have considerable existing metagenomic (from viruses) as well as rRNA sequence data. We used these local databases, including all contigs from our previously published 5-year viral metagenomic dataset^13^, SILVA v. 132^15^ and our collection of rRNA clones collected as part of our long-term SPOT ecological time series^16^. We updated the characterization of viral contigs in our 0.02-0.2 μm size fraction viral metagenome, by adding newer and more sensitive virus-finding tools to the previous application of Virsorter and VirFinder (confirmatory), specifically DeepVirFinder, CheckV, MEGAN-LR, VirSorter, and homologies to the Tara Ocean proteomic datasets (see methods). We then mapped reads with an overlap > 100 bp and > 95% identity to all the assembly from San Pedro Virome dataset, Searching for rRNA ligated to newly identified viral contigs, we identified 699 mRNA-rRNA chimeric reads, which represented 46 different viral contigs linked to 16S or 23S rRNA (Figure 2a, Suppl. File 1). We found more associations between mRNA and 16S than 23S rRNA (Fig 2a), despite 23S being longer, perhaps suggesting 16S has more accessible loops to allow enzymatic cleavage and re-ligations.

**Figure 2.**
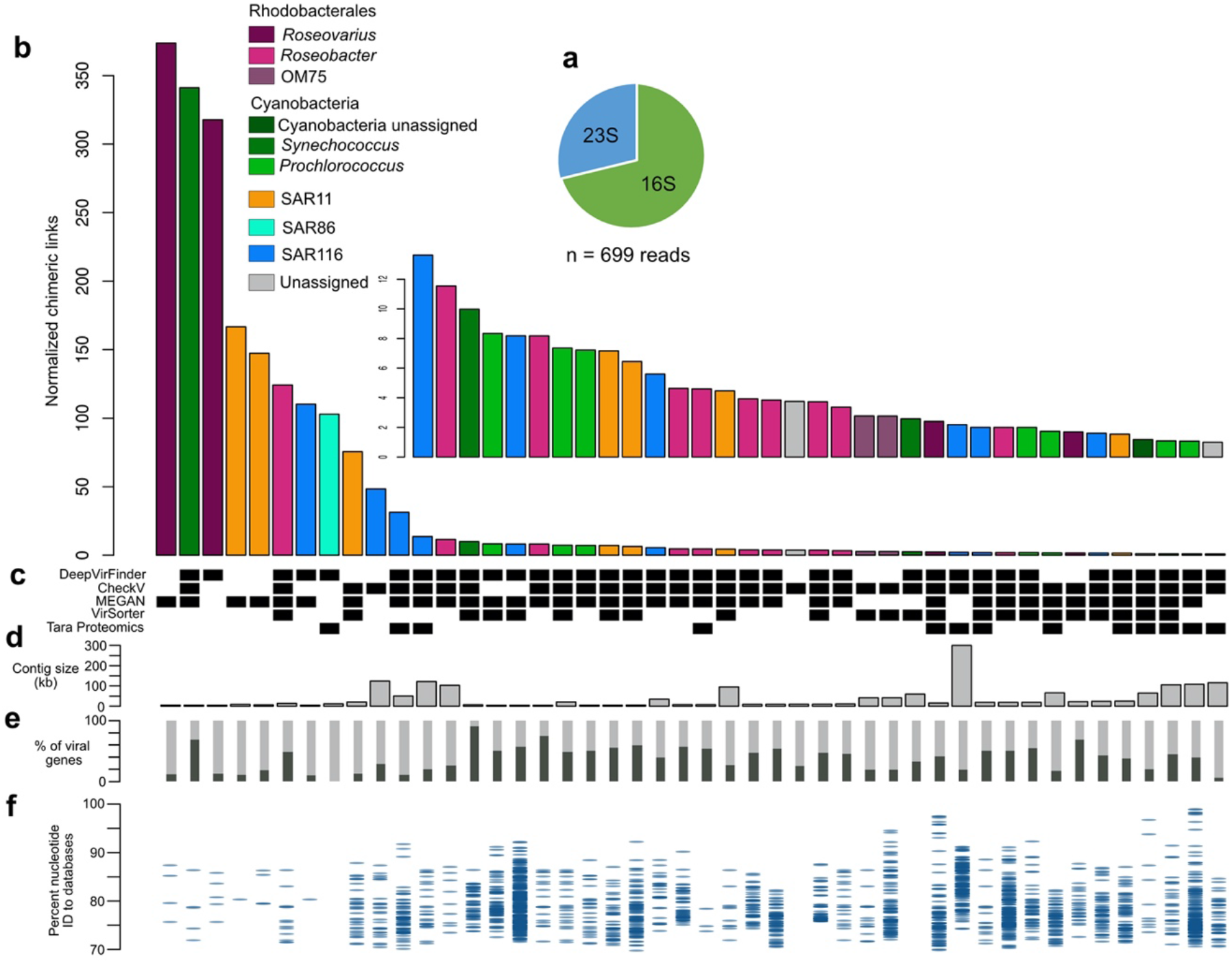
Novel virus-host associations discovered by RNA proximity ligation. **a)** More rRNA-mRNA chimeras are formed with 16S than 23S rRNA. **b)** Plot of the viral contigs (each bar a different contig) ranked from most to least normalized number of chimeric reads, with colors reflecting the taxonomy of the host from the rRNA within the chimeras. **c)** Black boxes indicating how a contig was identified as viral, methods named on the left. **d)** Length in kb for each contig. **e)** Percentage of genes (dark gray) within each contig currently identified as viral (note uncultivated virus genes are often not known as such) **f)** Distribution of similarity between gene matches obtained from within each contig and genes in public environmental virus databases (GOV and viral isolates from NCBI).

Our 46 identified viruses include those to hosts for which viruses were previously unknown, significantly the first SAR86 virus (Figure 2b, Suppl. File 1), particularly notable because this gammaproteobacterial group is globally abundant in seawater^19^. This virus appears to be an abundant member of the community at the transcriptomic level, though surprisingly it has no significant hits in the global ocean virome (GOV) dataset (Figure 2f), suggesting it may be regional or ephemeral. We also describe the first putative OM75 (alphaproteobacterial) viruses, and abundant Roseobacter phages unlike those previously reported (Figure 2; Suppl. Table1). Cyanobacterial viruses were well represented in the host-virus chimeras characterized by our methods, as expected due to their abundance in our samples, as well as their prevalence in the cultivated virus database. Phylogenetic 16S assignment divided them between *Prochlorococcus* (N= 7) and *Synechococcus* (N=3), and one whose associated 16S fragments did not allow us to distinguish between those two genera (Suppl. File 1; Figure 2b-f). We also found abundant phages associated with various Alphaproteobacteria, divided among Rhodobacteraceae (N= 16), SAR116 (N=8) and phages infecting the abundant SAR11 (N=8). Some of these host assignments, due to the abundance of the chimeras (many 16S hits), can be placed to the exact amplicon sequence variant (ASV) level, such as a *Roseovarius*, with multiple viral contig links to a single 16S ASV (Suppl. Table 1, Figure 3, below), suggesting that they are probably fragments of the same viral genome.

**Figure 3.**
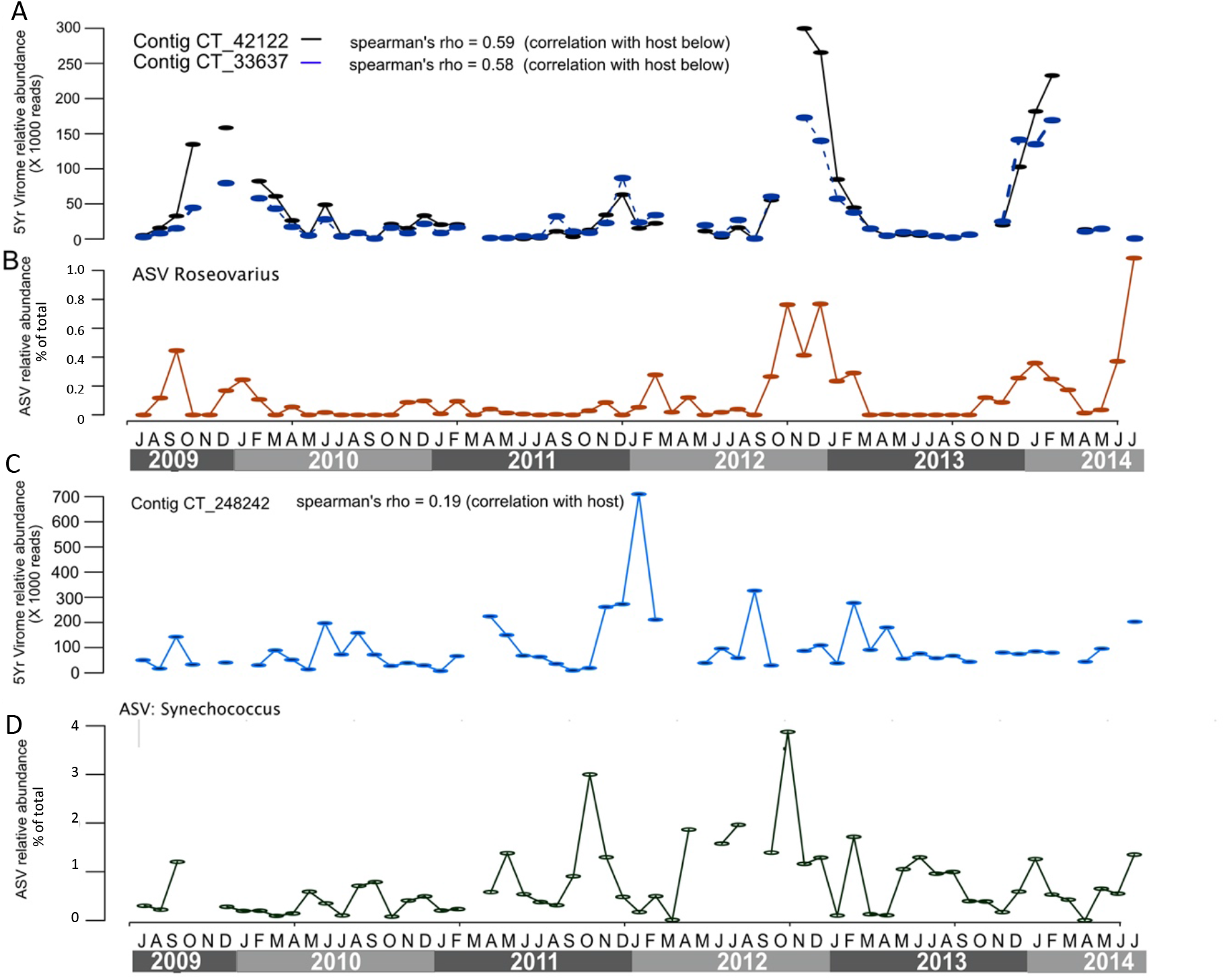
Tracking putative virus-host pairs over five years of ~monthly sampling at the San Pedro Ocean Time Series. Data are from 16S rRNA amplicons (host ASVs, are shown as percentage of the total community) and normalized recruitment of 0.02-0.2 μm (viral) size fraction metagenomic reads to viral contigs. **a)** Two virus contigs that both match the same putative *Roseovarius* host which is shown in **b.** These two contigs are both short (6.1 and 5.4 kbp) and considering the same host match and nearly identical dynamics of both contigs, are probably from the same virus. Note the general correspondence in abundances over time of contigs and presumed host, suggesting the virus largely follows its host abundance on this monthly time scale. **c)** and **d)** show an abundant viral contig and its presumed *Synechococcus* host, but in this case there is little correspondence between the dynamics, possibly due to strain variations we cannot detect by 16S and short read recruitment alone.

Due to the constraints of the bioinformatic methods used to identified viral contigs (complete dependencies on databases), it is difficult to identify fully novel viruses, and it is possible that linkages to many truly novel viruses have been missed by our conservative approach. We assessed the novelty of the viral lineages linked to particular hosts in our analysis, by identifying the percentage of known viral genes within a contig (Figure 2e) and the distribution of nucleotide identities to sequences from GenBank and metagenomic projects (Figure 2f)^14,20^. We see that most of the contigs have either a good percentage of hits that can be identified as viral or are represented in public metagenomic projects. Exceptions include the novel SAR86 virus (mentioned above), an unassigned cyanophage and a phage that we couldn’t place to any phylogenetic group due to the scarcity of the chimeric reads, which appear to be poorly represented in metagenomic assemblies. Beyond absolute novelty, our experiments revealed viral groups previously reported only to infect very different host groups than we report here, for example a novel T7-like Roseobacter phage (CT18917, Rank 6, Figure 2) with distant hits to T7 cyanobacterial phage (Suppl. File 1), and novel SAR11 viruses (CT_SN_17500 and CT_SN_38734, Ranks 20 and 21, Figure 2) that appear distantly related to enterobacterial T5 phages. This expands the range of known viruses infecting this numerically dominant ocean clade.

We can track viruses and their presumed host abundances from our San Pedro Time series, and interestingly we find contrasting virus-host patterns. The tracking is possible because we used the assemblies from our recently completed five-year viral metagenomic survey as a database to find viral sequences, and we can estimate relative abundances via read recruitment to those data. We also have time series data on potential host relative abundances from amplicon sequencing of SSU rRNA. Some of the linked viral contigs have many chimeric reads, and this allowed us to pin-point the specific 16S ASV associated to the host infected by this virus. So we tracked the long-term (5Yr) virus-host dynamics of these associations (Fig 3). Here we show three cases, one with a match to a *Synechococcus* ASV, and two with perfect matches a *Roseovarius* ASV (Suppl. File 1). The abundance of the host ASV in the latter case across time closely matches the dynamics of the virus in this case (Fig 3), consistent with a persistent virus infection where the virus essentially tracks the dynamic host abundances over many months. Yet that is not the only pattern we observed; we also were able to find the specific host for the third most abundant contig in the 5-year virome, and this cyanophage (Figure 3c-d) and its associated *Synechococcus* ASV are both dynamic but do not closely track each other. Perhaps this is due to strain-level variations in viruses and/or hosts that control the extent of infection, yet variation cannot be detected by short read recruitment nor fairly conserved 16S sequences; such strain variation is part of the Red Queen-like dynamics we previously reported for this location^13^. Similar *Synechococcus* strain variation in apparent infection dynamics was also reported for this location by Ahlgren et al^21^. Note also the cyanophage virus-host pair are both much more abundant than the *Roseovarius* pair, which may also relate to the difference in patterns.

We recognize there are potential shortcomings in requiring the virus has a well-assembled contig in order to match to a host; we know due to high genomic variability, it is often difficult to assemble many viral contigs in the first place, especially from only one or a few samples^13,22,23^. So as an alternative approach avoiding the need for assemblies, we searched for chimeric sequencing reads that aligned to known virus marker genes. We found 171 reads that match both cyanomyoviral marker gp20 or myoviral marker gp23 as well as a 16S rRNA for host phylogenetic placement (Figure S2). Not surprisingly, these were enriched for cyanobacterial viruses (which has a large cultured database) but they also matched, SAR11, SAR92, OM162 and Puniceicoccales (Verrucomicrobia), groups for which we have relatively few, if any, previously known viruses. This shows that our method can operate at the read level, and although the information is not as satisfactory as having a long viral contig or genome, it is still valuable to know a particular host is infected by a virus for which we now have at least one specific marker that can be tracked in metagenomes (for occurrence) or metatranscriptomes (for active infections) via read recruitment.

In comparison to other methods that aim to link unknown viruses and hosts, this approach has some obvious advantages. It is much more specific than k-mer baser methods^3^, which are general purpose and provide probabilities of matches, but typically do not narrow the hosts down to better than genus or family levels with confidence. It is more high-throughput, and less costly per match, than methods requiring sequencing sorted cells after amplification^5,6^. It is most similar to DNA-DNA proximity ligation methods^8,9^, and the principal differences are that (1) our approach catches the virus in the act of transcription while DNA-DNA approaches will link any DNA within in close proximity, perhaps catching non-infection situations or unsuccessful infections, and (2) we can place any host on a phylogenetic tree (or find an exact match if available) by its 16S (or 23S) rRNA sequence while the general DNA-DNA proximity ligation method requires host genome sequence information to identify it, and such information on most naturally occurring organisms is limited. One important limitation of our method is that by the nature of sequencing, the information in short reads is limited. We expect that future developments such adapting long read sequencing^24^ will help overcome this shortcoming.

In conclusion, we have adapted molecular biology technique based on proximity ligation and applied to a first field sample to uncover novel virus-host associations. We anticipate our methods will be widely applied and improved upon to study the dynamics of interaction networks in natural environments. Finally, because the proximity ligation is non-specific for viruses and in fact can associate any translated protein with the ribosome doing the translation, it can link environmental functions to taxonomic units, much needed for a mechanistic modeling of a changing ocean.

---

**Suppl. Figure 1.**
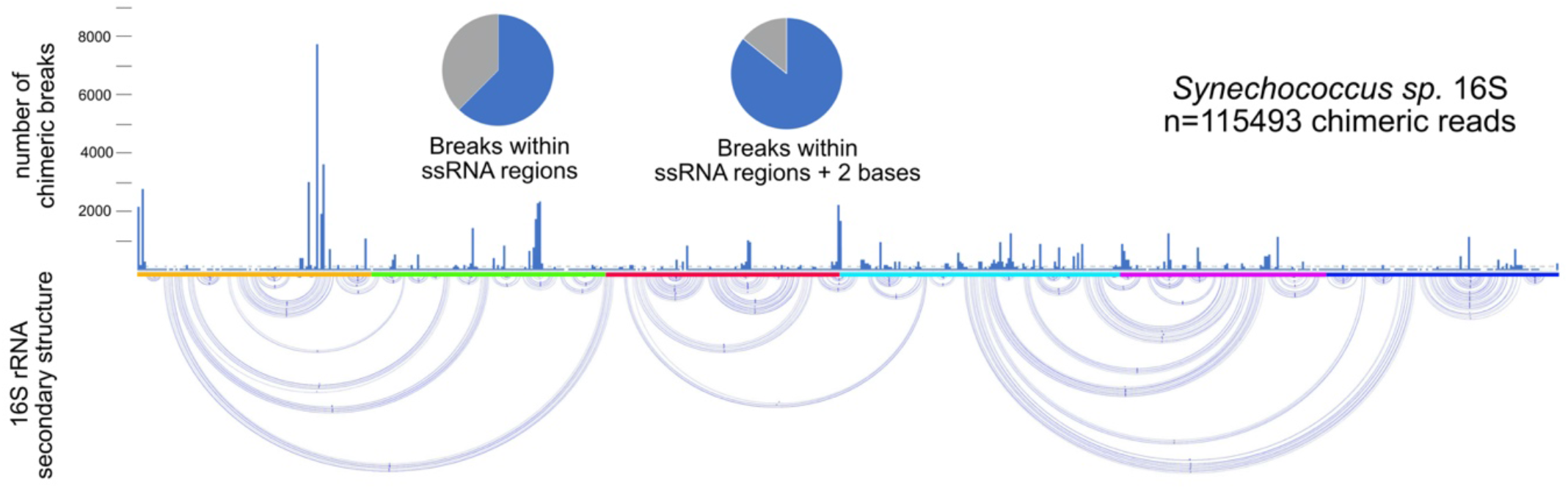
In accordance with the premise of our experiment the vast majority of the chimeric break points occur in single stranded (loop) regions. Bottom panel of the figure represents the known secondary structure of the 16S ribosomal RNA from *Synechococcus* (https://crw-site.chemistry.gatech.edu/RNA/Structures/d.16.b.Synechococcus.sp.pdf). Top panel shows the number of chimeric read breakpoints at each base, scale on the left side. Pie charts show the majority of the breaks were at single stranded regions, i.e. loops (or within 2 bases to accommodate natural variation in an environmental sample). Single stranded regions represent only 1/3 of the length of the 16S rRNA yet it accounts for up to 85% of the breakpoints; note the breaks are relatively non-random and appear to be enriched at the beginning of the sequence.

**Suppl. Figure 2.**
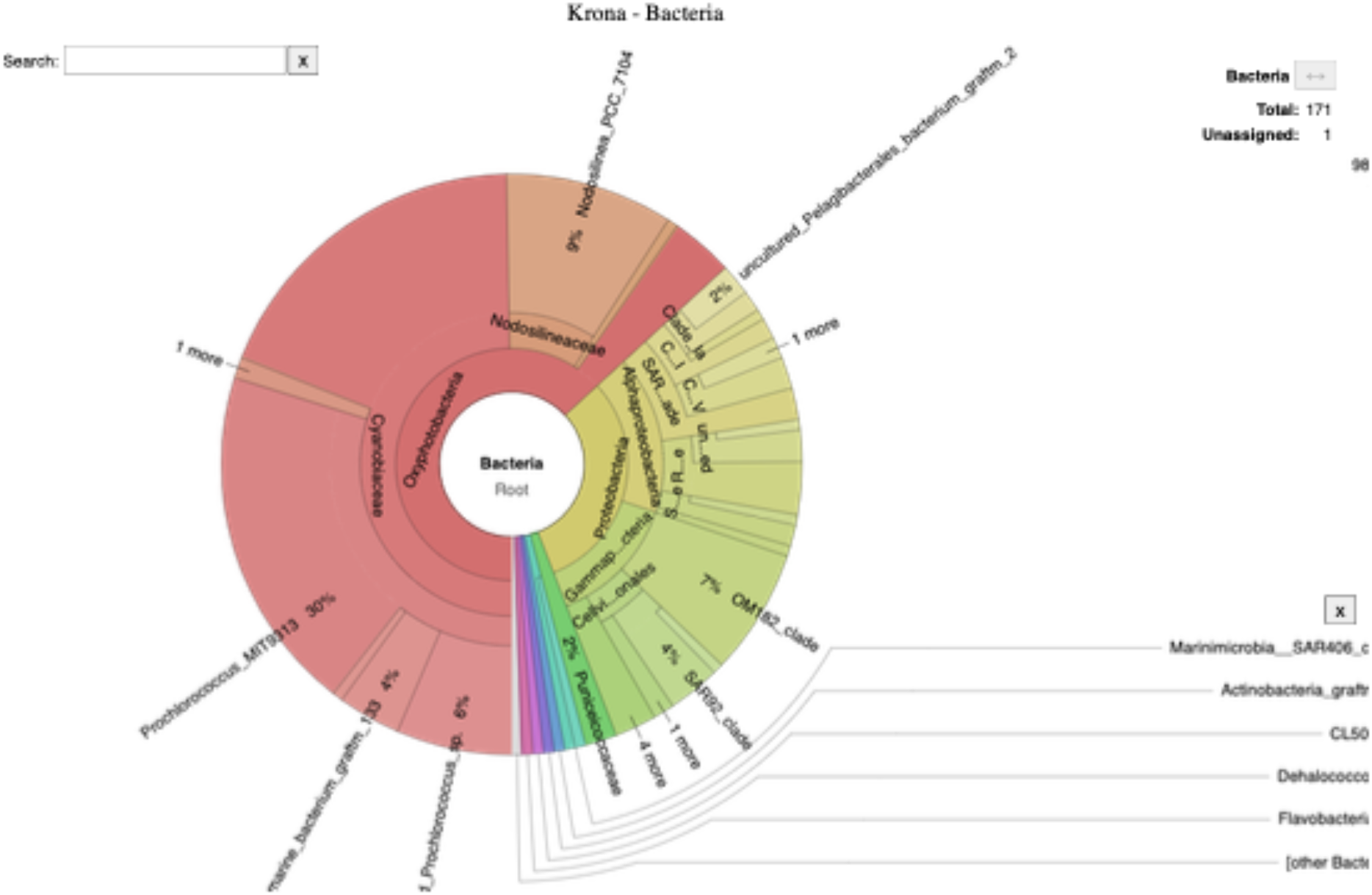
Read level analysis reveals potential links between T4-like viral genes and diverse hosts. We identified reads that matched T4 like genes, and the 16S region of the rRNA-mRNA chimera was then placed onto a phylogenetic tree using the graftm^25^ 16S package. Not unexpectedly (due to their high representation in the database and also high natural abundance), cyanobacteria accounted for 60 % of the chimeric reads, but nonetheless T4 is a widespread family with a great variety of marine hosts.

## Materials and Methods

### Sample collection

Seawater was collected aseptically using a bucket previously stored in 5% HCl during the February 2020 cruise of the San Pedro Ocean Time series (https://dornsife.usc.edu/spot/); 16 L were filtered through a 1 um fiber glass AE filter (Millipore) to remove larger particles, as well as larger phyto- and bacterioplankton. Filtrate was then filtered unto a 0.22 um Sterivex cartridge (Millipore, with a Durapore filter); filter was then dried by pushing air with a 50 mL syringe. We immediately proceeded to crosslink our sample on this sterivex filter.

### Crosslinking and RNA extraction

Samples were fixed with 1% formalin (0.4% formaldehyde) to create cross-links between adjacent proteins, 2 mL of formalin were added inside the sterivex cartridges, filter ends covered with locking luer caps, and filter tilted gently over 5 mins. Excess liquid was pushed out of the sterivex using a clean syringe. Formalin was then quenched by filling the sterivex cartridge with 250 nM glycine for 20 minutes. Glycine was pushed out using a clean syringe. Extraction of RNA-Protein complexes was accomplished largely following the directions described by Trendel et al.^12^ modified to accommodate a filter enclosed in a sterivex cartridge. 1.5 ml of Trizol (Sigma-Aldrich) were added inside the cartridge, which was capped and mixed for 5 minutes on a vortexer; sterivex cartridge was then opened again and 0.3 mL of chloroform were added to induce phase separation. The contents of the sterivex filter (~1.8 mL) were the transferred to a LoBind 2mL tube and rested at room temperature for 5 minutes. Tubes were centrifuged at 7000 x g and 4C for 10 minutes. RNA was recovered from the aqueous part following the directions of manufacturer. RNA-Protein complexes were recovered from the interface as recommended by Trendel et al^12^, resuspended in 100 uL of DEPC water. RNA from interface and aqueous fraction was then subjected to two rounds DNase I (NEB) treatment, 100 uL of DNase I (100 Units) and 1.8 mL of 10X DNase buffer to a final reaction of 2mL. Each time the RNA was concentrated and cleaned by one round of isopropanol and then one of ethanol precipitation. It was suspended in 100 uL in DPEC water, quantified with Qbit RNA, we immediately proceeded to the following steps.

### RNA cleavage and ligation

RNA was randomly cleaved generating accessible ends both in the rRNA and mRNA, largely based on the methods by Sharma et al.^11^ Multiple 20 uL reactions were run in parallel, 100 ng of RNA (RNA-protein complexes) each, with 2 Units of S1 enzyme (ThermoFisher, 2 uL of a 1:100 dilution) and 1X S1 Buffer. Reactions were run for 30 mins at room temperature, reactions were stopped by phenol chloroform extraction. Free mRNA and rRNA ends were then ligated into circular forms with a circRNA ligase (Lucigen); it favors ligation events between proximal RNA ends^11^ and it has limited activity at high temperature, which would favor ligation of proximal ends only. Specifically, multiple 20 uL reactions were run in parallel, 50 ng of S1-digested RNA was incubated with 2 uL of 10X circRNA ligase buffer at 85C for 2 minutes. Tubes were transferred to ice where 1 uL of 10 mM ATP and 1 uL of circRNA ligase were added. Reactions were run at 60C for 60 minutes.

### Enrichment for Chimeric reads, crosslinking reversal

Multiple 25 uL reactions were run in parallel, reactions were set up by adding 0.5 uL of RNaseR (Lucigen), 2.5 uL of RNase R buffer and 2 uL of water to the previous set ups (ligated RNA). These reactions were incubated for 10 minutes at 37C. RNase reactions were stopped with Proteinase K by adding 30 uL of 2X Proteinase K buffer, 3 uL of Proteinase K and 2 uL of water, reactions were incubated 30 minutes at 60C. Proteinase K treatment also reverses the crosslinking. RNA was purified by phenol chloroform extraction and suspended in nuclease free water.

### Retro transcription and library generation

Multiple 20 uL reactions were run each with 100 ng of RNA using the NEB first strand synthesis module (Parts no. E7525). 13.5 uL of RNA and 1 uL of random primers were incubated at 65C for 5 minutes then put on ice. To this, 4 uL of reaction buffer and 0.5 uL of RNase inhibitor were added, incubated at 25 C for 2 minutes. Finally 1 uL of Protoscript II reverse transcriptase was added and incubated 10 min at 25 C, 50 at 42C and 15 min at 70C, then put immediately on ice. We then proceeded with the NEB Second Strand Synthesis Module (Parts no. E7550), by adding 8 uL of the dNTP mix, 4uL of the enzyme mix and 48 uL of nuclease-free water to a total volume of 80 uL. Reactions were incubated at 16C for 1 hour. Samples were pooled, cleaned and concentrated using a 1.2 X AMPure magnetic beads (Beckman-Coulter) and resuspended in 40 uL of low EDTA TE. Sequencing libraries were generated using the Ovation Ultralow V2 (NuGen) with 14 amplification cycles. Libraries were sent out for sequencing on a 2X250 PE NovaSeq, with a final sequencing depth of 41 M (aqueous fraction) and 35 M (interface) on each sample. Data from these two samples was pooled and treated identically in subsequent steps.

## Bioinformatic analysis

### Quality Control and read merging

Reads were qc’ed using fastp^26^ using a minimum quality score of 15 covering at least 75 % of the read length (Options: “-q 15-u 25”); we allowed for a relatively low score value since we use the reads for read recruitment. Reads were then merged using fastq-join^27^ allowing a maximum 10% differences on a minimum 20 bases overlap (Options: “-p 10 -m 20”). In practice this generated an insert size range from 250 bp to 480 bp.

### Custom perl scripts

Custom perl scripts that were written to parse and analyze the data from our experiment have been deposited at https://github.com/phagenomics/VirHostLinker, they are referenced throughout the methods as <script>.pl<input files>.

### A posteriori evaluation of crosslinking specificity

The initial dataset (76M reads) was blasted against the collection of single cell genomes from Pachiadaki et al.^17^ with all default options (Options= “blastn -outfmt 6 – num_threads 8”), the 16S sequences from these SAGs were clipped out prior to running the blast program. We then extracted a list of sequences that matched anything within that database with a percent identity higher than 95% over at least 100 bp using a linux one liner (“awk ‘($3 >= 95)’ | awl ‘($4 >=100)’ | awk ‘{print $1}’ | uniq > LIST”). We then extracted all these reads from the initial dataset using custom perl scripts (perl splitRNA.pl LIST). This subset of sequences was then blasted against all the previously extracted 16S sequences from these SAGs (N = 4726 sequences), hits with a percent identity higher or equal to 98% over at least 100 bp were chosen selected as high quality chimeric reads (“awk ‘($3 >= 98)’ | awl ‘($4 >=100)’ | awk ‘{print $1}’ | uniq” as above). Only the top blast hit of the 16S region is considered, while all the hits higher than 95% ID on the non-rRNA regions were considered equally good. This last constraint implies that if a read matches one 16S sequence and five non-16S SAG sequences it will appears counted in the matrix of figure 1b five times. We only validated using the 16S/18S gene.

### Identification of novel-virus host linkages

The initial dataset (76M reads) was blasted against the final viral contig assembly from our previous work^13^ (N = 99907 contigs, > 5Kb) with all default settings (Options= “blastn -outfmt 6 – num_threads 8”). We then extracted a list of sequences that matched anything within that database using a Linux one liner (“awk ‘{print $1}’ | uniq”), at this point we did not filter for any level of identity. We then extracted all these reads from the initial dataset using custom perl scripts (“perl splitRNA.pl LIST”). This subset of sequences was then blasted against SILVA^15^ and an in house collection of local near-full length 16S-ITS clone sequences^18^. High quality chimeric reads were then identified using custom Perl scripts, and were chosen if between 40 and 60 percent of the read is covered by a match in one database and the rest by another match in the other database and if the minimum length of either alignment is 100 bp (“perl PartialAlignment.pl”). We found 1.5 M reads that were identified as high-quality chimeras linking contigs from the 5Yr virome and a ribosomal RNA molecule. Many of these hits were to cellular fragments within the 5Yr virome, so we curated futher using different bioinformatic pipelines. We narrowed the final contig list to include only contigs that met one of the following criteria: VirSorter^28^ (Categories 1 to 3), DeepVirFinder (Scores > 0.9) were selected, MEGAN-LR^29^, CheckV^30^ and by finding homologies to proteins within Tara viral proteomics datasets^31^ (see below); The identification tool used for each contig is depicted in figure 2C, and all the values from each pipeline are in Suppl. Table 1. Each read was uniquely assigned to the contig as the top hit with the additional minimum identity to be 95%. The other end of the reads was assigned to the top hit in the previously described clone database and to silva if the closest match within the local clones databases was lower than 95% and/or the chimeras was formed with 23S rRNA (Suppl. File 1).

### Additional curation of viral contigs

Contigs were annotated using the top blastp hit to nr (Accessed August 2020). Additionally, we used the virus proteomic^31^ dataset from Tara to inform some of the annotations of our dataset as viral, this second approach identified structural proteins in 33 contigs (Suppl. File Table 1). For MEGAN-LR, contigs were aligned using LAST^32^ to the NCBI nt database (June 20, 2019) and the results were input to MEGAN-LR^29^ using the lowest common ancestor (LCA) algorithm. CheckV was run using default settings using checkv-db-v0.6. Viral contigs were predicted de novo on contigs longer than 2000 bp using DeepVirFinder^33^ requiring a p-value of 0.01.

### 16S PCR amplification and ASV calling

Prokaryotic DNA (0.2-1 um fraction) corresponding to the sampling dates overlapping with our metagenomic work was extracted as described by Chow et al.^34^. V4 and V5 regions were amplified using the primers described by Parada et al.^35^, following the methods described by Yeh et al^36^. ASVs used in this study have been deposited at https://github.com/phagenomics/VirHostLinker.

## Code Availability and Supplementary Information

Custom code and supplementary files (for pre-print version) available at https://github.com/phagenomics/VirHostLinker. Formatted bash scripts to obtain the results presented in this manuscript can also be found there.

## Data availability

All data needed to evaluate the conclusions in the paper are present in the paper or the supplementary materials. Final cross-assembled sequences and raw sequencing data are deposited at NCBI under the BioProject PRJNA672948.

## Author Contributions

JCIE and JF conceived and designed the experiment; SL, YCY, SH, and AML contributed with bioinformatic pipelines, analyses, and databases. JW contributed with data visualization and statistical analyses. JCIE, SL, JW, YCY, SH, AML and JF contributed and commented on the final form of this manuscript.

